# The Genetic cost of Neanderthal introgression

**DOI:** 10.1101/030387

**Authors:** Kelley Harris, Rasmus Nielsen

## Abstract

Approximately 2-4% of genetic material in human populations outside Africa is derived from Neanderthals who interbred with anatomically modern humans. Recent studies have shown that this Neanderthal DNA is depleted around functional genomic regions; this has been suggested to be a consequence of harmful epistatic interactions between human and Neanderthal alleles. However, using published estimates of Neanderthal inbreeding and the distribution of mutational fitness effects, we infer that Neanderthals had at least 40% lower fitness than humans on average; this increased load predicts the reduction in Neanderthal introgression around genes without the need to invoke epistasis. We also predict a residual Neanderthal mutational load in non-Africans, leading to a fitness reduction of at least 0.5%. This effect of Neanderthal admixture has been left out of previous debate on mutation load differences between Africans and non-Africans. We also show that if many deleterious mutations are recessive, the Neanderthal admixture fraction could increase over time due to the protective effect of Neanderthal haplotypes against deleterious alleles that arose recently in the human population. This might partially explain why so many organisms retain gene flow from other species and appear to derive adaptive benefits from introgression.

## 1 Introduction

In recent years, prodigious technological advances have enabled extraction of DNA from the remains of our extinct Neanderthal relatives [1]. Analysis of this ancient DNA revealed that Neanderthals had lower genetic diversity than any living human population [2, 3]. By analyzing patterns of divergence between distinct Neanderthal haplotypes, Priifer et al. inferred that Neanderthals experienced a strong population bottleneck, lasting approximately ten times longer than the out-of-Africa bottleneck [2, 4, 5, 6].

A classical consequence of population bottlenecks is that they interfere with natural selection by increasing evolutionary stochasticity [7, 8]. When effective population size is small and genetic drift is therefore strong, weakly deleterious alleles have a tendency to persist in the population as if they were neutral. Neanderthal exome sequencing has confirmed this prediction, providing direct evidence that purifying selection was weaker in Neanderthals than in humans [3, 9]. Compared to humans, Neanderthals have a relatively high ratio of nonsynonymous to synonymous variation within proteins, indicating that they probably accumulated deleterious nonsynonymous variation at a faster rate than modern humans do.

It is an open question whether archaic hominins’ deleterious mutation load contributed to their decline and extinction. However, there is clear evidence that Neanderthals escaped total genetic extinction by interbreeding with the anatomically modern humans who left Africa between 50 and 100 thousand years ago [1]. In Europeans and Asians, haplotypes of Neanderthal origin have been inferred to comprise 2-4% of each individual’s genome. When pooled together, these Neanderthal haplotypes collectively span about 30% of the human reference sequence [10, 11].

The introgression of Neanderthal alleles related to hair, skin pigmentation, and immunity appear to have provided non-Africans with adaptive benefits, perhaps because Neanderthals had preadapted to life in Europe for thousands of years before humans left Africa [10, 11, 12, 13, 14]. However, these positively selected genes represent a tiny fraction of Neanderthal introgression’s genetic legacy. A larger number of Neanderthal alleles appear to have deleterious fitness effects, with putative links to various diseases as measured by genome-wide association studies [10, 15].

The distribution of deleterious mutations in humans has been the subject of much recent research. A controversial question is whether the out-of-Africa bottleneck created differences in genetic load between modern human populations [16, 17]. Some previous studies concluded that this bottleneck saddled non-Africans with potentially damaging genetic variants that could affect disease incidence across the globe today [18, 19, 20], while other studies have concluded that there is little difference in genetic load between Africans and non-Africans [21, 9]. Although previous studies have devoted considerable attention to simulating the accumulation of deleterious mutations during the out-of-Africa bottleneck, none to our knowledge have incorporated the fitness effects of introgression from Neanderthals into non-Africans.

In this paper, we quantify the deleterious effects on humans of introgression with Neanderthals with a high mutational load. We present simulations showing that archaic introgression may have had fitness effects comparable to the out-of-Africa bottleneck, saddling non-Africans with weakly deleterious alleles that accumulated as nearly neutral variants in Neanderthals.

## 2 Results

To assess the fitness effects of Neanderthal introgression on a genome-wide scale, we used forward-time simulations incorporating linkage, exome architecture, and population size changes to model the flux of deleterious mutations across hominin species boundaries. We describe three main consequences of this flux, which are not mutually exclusive and whose relative magnitudes depend on evolutionary parameters such as the distribution of dominance coefficients and fitness effects of new mutations. One consequence is strong selection against early human/Neanderthal hybrids, implying that the initial contribution of Neanderthals to the human gene pool may have been much higher than the contribution that persists today. A second consequence is depletion of Neanderthal ancestry from conserved regions of the genome, a pattern that has been previously inferred from genetic data [10, 11] and interpreted as evidence for partial reproductive incompatibilities between humans and Neanderthals. A third consequence is the persistence of deleterious alleles in present-day humans, creating a difference in mutation load between non-Africans (who experienced Neanderthal admixture) and Africans who did not.

### 2.1 The Reduced Fitness of Neanderthals

Our first step toward quantifying these three consequences of introgression was to estimate pre-admixture mutation loads in humans and Neanderthals. We accomplished this using simulations where all humans and Neanderthals experience deleterious mutations drawn from the same distribution of fitness effects (DFE), such that any differences in mutation load are driven by differences in demographic history. Because the fitness effects of noncoding mutations are difficult to measure, we restricted our attention to deleterious mutations that alter protein coding sequences (nonsynonymous or NS mutations). There have been several estimates of the distribution of selection coefficients in human protein coding genes [22, 23, 24, 25]. We here use the estimates of Eyre-Walker, *et al*. who found that the DFE of human NS mutations is gamma-distributed with shape parameter 0.23 and mean selection coefficient -0.043 [26]. Although it is probably unrealistic to neglect the fitness effects of synonymous and non-exonic mutations, it is also conservative in that additional deleterious mutations would only increase the human/Neanderthal load difference beyond the levels estimated here. Although little is known about the mutational DFE outside coding regions, any deleterious mutations that occur will fix in Neanderthals with higher probability than in humans; in addition, any beneficial mutations that occur will fix with higher probability in humans than in Neanderthals.

Using the UCSC map of exons from the hg19 reference genome, we assume that each exon accumulates NS mutations with fitness effects sampled uniformly at random from the distribution estimated by Eyre-Walker et al. Since the human germline mutation rate is approximately 1.0n × 10^−8^ mutations per site per generation [27, 28] and approximately 1/3 of new mutations in coding regions should not change the amino acid sequence, we set the NS mutation rate to be 7.0 × 10^−9^ mutations per site per generation. No deleterious mutations occur between exons, but recombination does occur at a rate of 1.0 × 10^−8^ crossovers per site per generation. We implemented this genetic architecture within the simulation program SLiM [29] by using the recombination map feature built into the simulator. Specifically, for each pair of adjacent exons separated by a gap of *b* base pairs, we represent this gap as a single base pair with recombination rate *b* × 10^−8^ per generation. Similarly, each boundary between two chromosomes is encoded as a single base pair with a recombination rate of 0.5 crossovers per generation. We chose to focus on the dynamics of the 22 autosomes, neglecting the more complex evolutionary dynamics of the X and Y chromosomes.

We allowed the mutation spectrum of this exome to equilibrate in the ancestral human/ Neanderthal population by simulating an ancestral population of size 10,000 for 44,000 generations. After this mutation accumulation period, the ancestral population splits into a human population of size 10,000 plus a Neanderthal population of size 1,000. The Humans and Neanderthals then evolve in isolation from each other for 16,000 more generations (a divergence time of 400,000 470,0 years assuming a generation time between 25 and 29 years). To a first approximation, this is the history inferred by Prüfer, et al. from the Altai Neanderthal genome using the Pairwise Sequentially Markov Coalescent [2]. Throughout, we assume log-additive interactions among loci. In other words, the fitness of each simulated individual can be obtained by adding up the selection coefficients at all sites to obtain a sum *S* and calculating the fitness to be exp(–*S*). The fitness of individual *A* relative to individual *B* is the ratio of their two fitnesses. For each of three different dominance assumptions, described below, three replicate simulations were performed and all results were averaged over the three replicates.

We ran three sets of simulations that differed in their assumptions regarding dominance coefficients of *de novo* mutations: one with fully additive effects, one with fully recessive effects, and one where mutations were partially recessive (all having dominance coefficient *h* = 0.1). We expect that the true distribution of dominance effects falls somewhere within the range of these extreme models. Although distributions of the dominance coefficient *h* have been inferred from viability data in mutation accumulation lines of yeast and *Drosophila*, these studies had limited power to classify weakly deleterious mutations with *s* < 0.01 [30, 31]. We have therefore avoided making assumptions about the distribution of dominance coefficients, instead describing the qualitative contrast between the effects of additive and recessive mutations. We use the same distribution of selection coefficients for the additive simulations and the recessive simulations to ensure that differences between their results are attributable to dominance effects alone. There is some evidence for an inverse correlation between *h* and *s* [30, 31, 32], meaning that weakly deleterious mutations are less often recessive than strongly deleterious mutations are. However, when Agrawal and Whitlock inferred a joint distribution of *h* and *s* from yeast data, they found that *h* is approximately gamma distributed given *s* [31], such that both additive and recessive mutations are expected to occur within each fitness class.

In our simulation with additive fitness effects, the median Neanderthal was found to have fitness 0.63 compared to the median human (Figure 1A). Assuming recessive fitness effects, the excess load accumulated by Neanderthals was even greater, with a median Neanderthal fitness of 0.39 compared to the median human ( Figure 1B). Such a large fitness disadvantage would have been incompatible with Neanderthal survival if they were competing with humans under conditions of reproductive isolation. In each case, the fitness differential was caused by accumulation of weakly deleterious mutations with selection coefficients ranging from 5 × 10^−5^ (nearly neutral in the larger human population) to 2 × 10^−3^ (nearly neutral in the smaller Neanderthal population). This agrees with asymptotic predictions that mutations with 2*Ns* > 1 are not affected by a bottleneck with minimum population size *N* [33].

**Figure 1:**
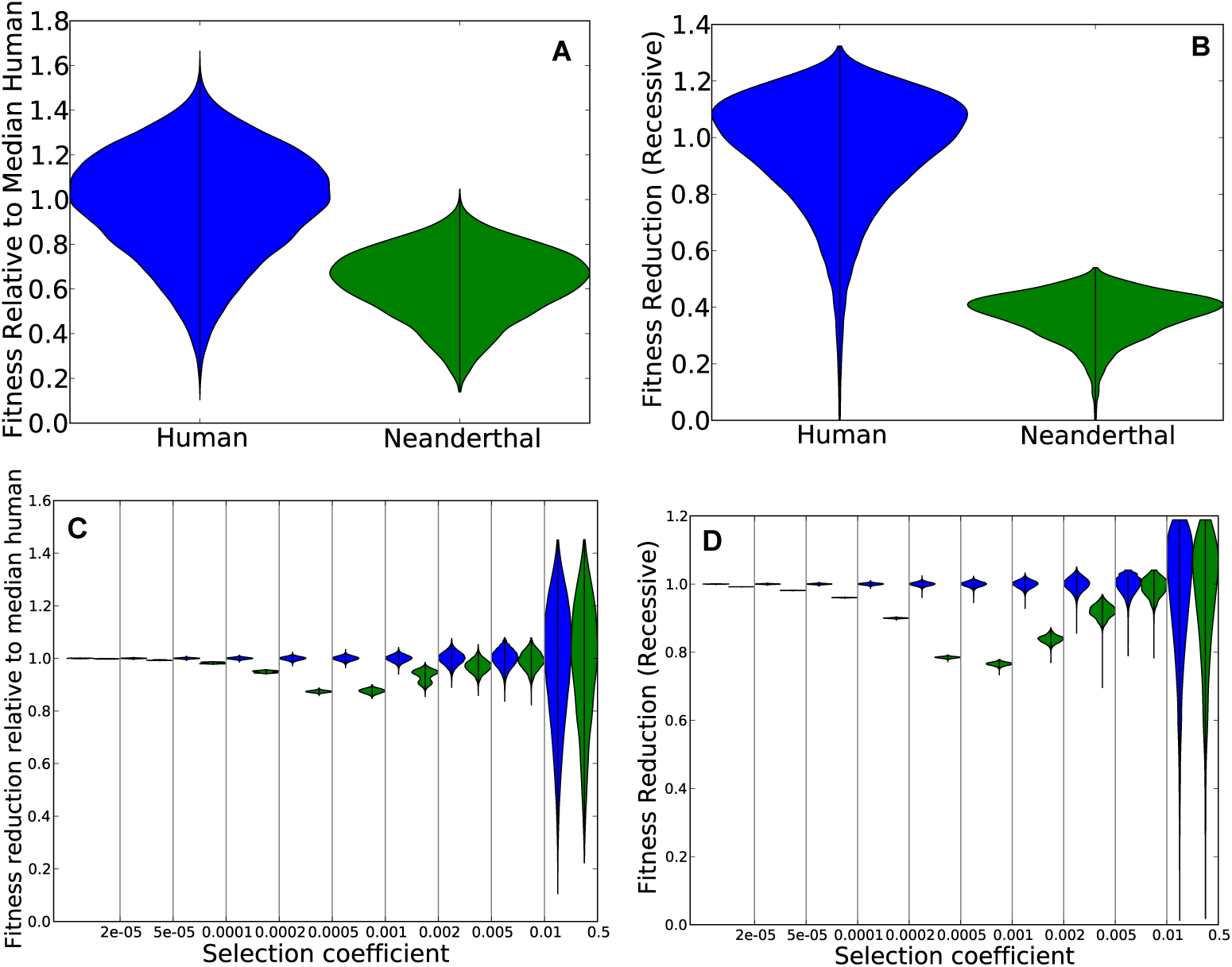
Panel A shows the distribution of fitness in Neanderthals versus non-admixed humans, assuming that their effective population sizes differed ten-fold since their divergence 16,000 generations ago, and assuming additive mutation effects. After simulating the two populations using SLiM, we calculated each individual’s fitness relative to the median human. The violin plots of the two distributions show significant variance within each population but lower fitness in Neanderthals. Panel B shows the same but for a model of recessive mutations. Panels C and D show the same data as in A and B, respectively, but stratified into different bins of selection coefficients. The fitness reduction due to mutations with *s* between 2 × 10^−4^ and 5 × 10^−4^, is much greater in Neanderthals than in humans. In contrast, the fitness reduction due to very weak effect mutations *s* < 2 × 10^−5^ is similar between the two populations, as is the fitness reduction due to strongly deleterious mutations with *s* > 0.005. This is expected as mutations outside this range are either effectively neutral in both populations or strongly deleterious in both populations.

To illustrate, we divided selection coefficient space into several disjoint intervals and measured how each interval contributed to the fitness reduction in Neanderthals. For each interval of selection coefficients 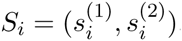, and each individual genome *G*, we calculated the mutation load summed across derived alleles with selection coefficients between 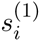 and 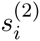 to obtain a load value *L_i_*(*G*). Given that *L_i_*(*G*_0_) is the median human load of mutations from the interval *S_i_*, the fitness reduction due to *S_i_*-mutations in a different individual *G*′ is exp(–(*L_i_*(*G*′) –*L_i_*(*G*_0_))). Figures 1C and D show the distribution of this fitness reduction. Variance between individuals is high for strongly deleterious mutations because an individual carrying one or two of these alleles is so much worse off than an individual who carries zero.

### 2.2 Recessive Mutations Lead to Positive Selection for Neanderthal DNA

We model Neanderthal gene flow as a discrete event associated with an admixture fraction *f*, sampling *Nf* Neanderthals and *N*(1 – *f*) humans from the gene pools summarized in Figure 1, and then allowing this admixed population to mate randomly for 2,000 additional generations. For each of the six simulated human and Neanderthal population replicates (three additive and three recessive) and each admixture fraction considered (*f* = 0, *f* = 0.01, and *f* = 0.1), we performed between 1 and 6 replicate introgression simulations that began with the same parent populations but randomly generated different human/Neanderthal hybrids. A Neanderthal gene flow date of 2,000 generations before the present is compatible with Fu, et al.’s claim that the admixture occurred 52,000-58,000 years ago [34], assuming a human generation time between 26 and 29 years. To simulate the out-of-Africa bottleneck, which affected humans around the time of admixture, we used a model based on the history inferred by Gravel, et al. from the site frequency spectrum of the 1000 Genomes data [5]. At the time of admixture (2,000 generations ago), the non-African population size drops from *N* = 10,000 to *N* = 1,861. 900 generations later, the size is further reduced to *N* = 1, 032 and begins exponentially growing at rate 0.38% per generation. We discretized this exponential growth such that the population size increases in a stepwise fashion every 100 generations (Figure 2). Because forward-time simulations involving large numbers of individuals are very time and memory intensive, we also capped the population size at *N* = 20,000 (the size that is achieved 300 generations before the present).

**Figure 2:**
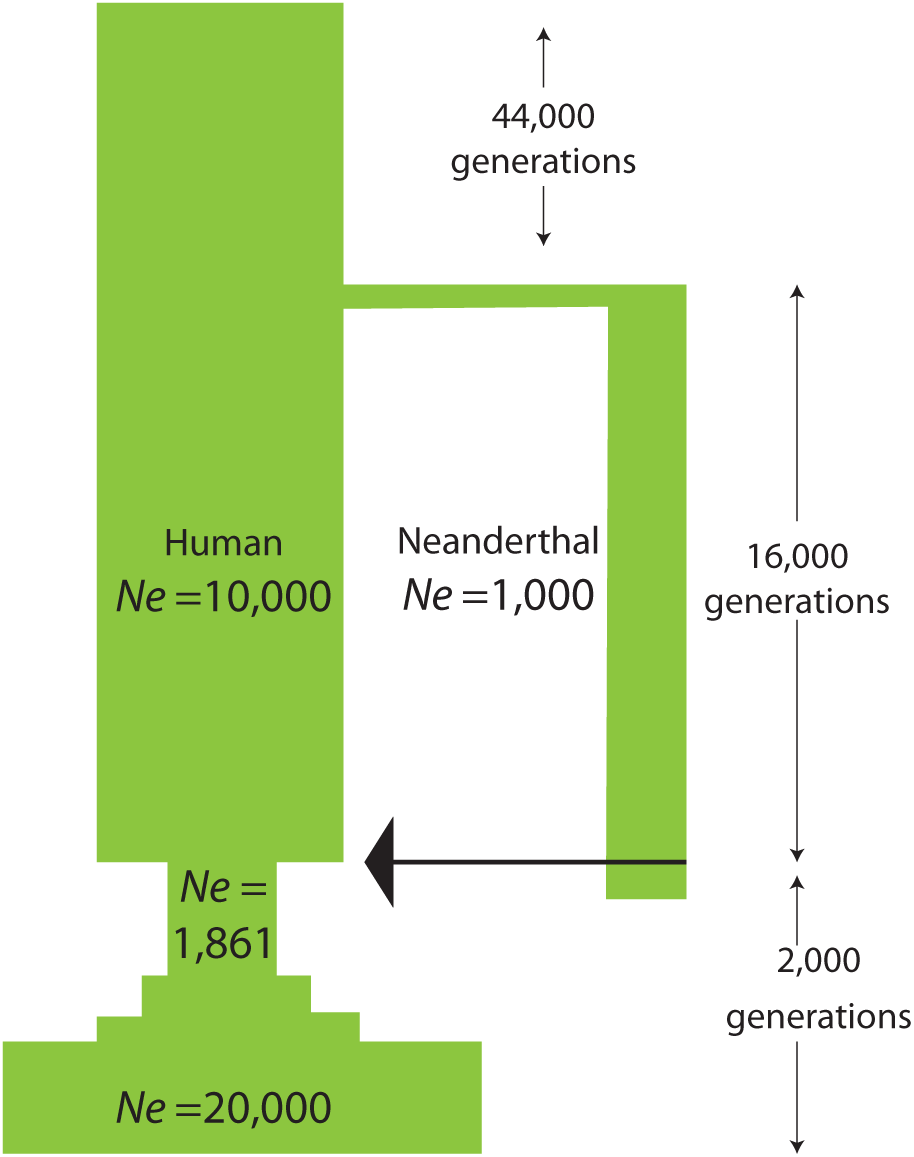
This cartoon shows the demographic history used to simulate Neanderthal and human genomes. A single ancestral population of size 10,000 is simulated for 44,000 generations to let the distribution of deleterious mutations reach equilibrium. At this point, a Neanderthal population of size 1,000 splits off. After 16,000 generations of isolation, Neanderthals and humans admix, followed by the out of Africa bottleneck and piecewise-constant exponential growth.

In the recessive-effects case, we found that the Neanderthal admixture fraction increased over time at a logarithmic rate (Figure 3). To quantify this change in admixture fraction, we added neutral marker mutations (one every 10^5^ base pairs) to the initial admixed population that were fixed in Neanderthals and absent from humans. The average allele frequency of these markers started out equal to the admixture fraction *f*, but was observed to increase over time. An initial admixture fraction of 1% was found to be consistent with a present-day admixture fraction around 3%, with most of the increase occurring over the first 500 generations. The selection favoring Neanderthal alleles is an example of dominance heterosis [35, 36, 37, 38], selection for foreign DNA that is protective against standing deleterious variation.

**Figure 3:**
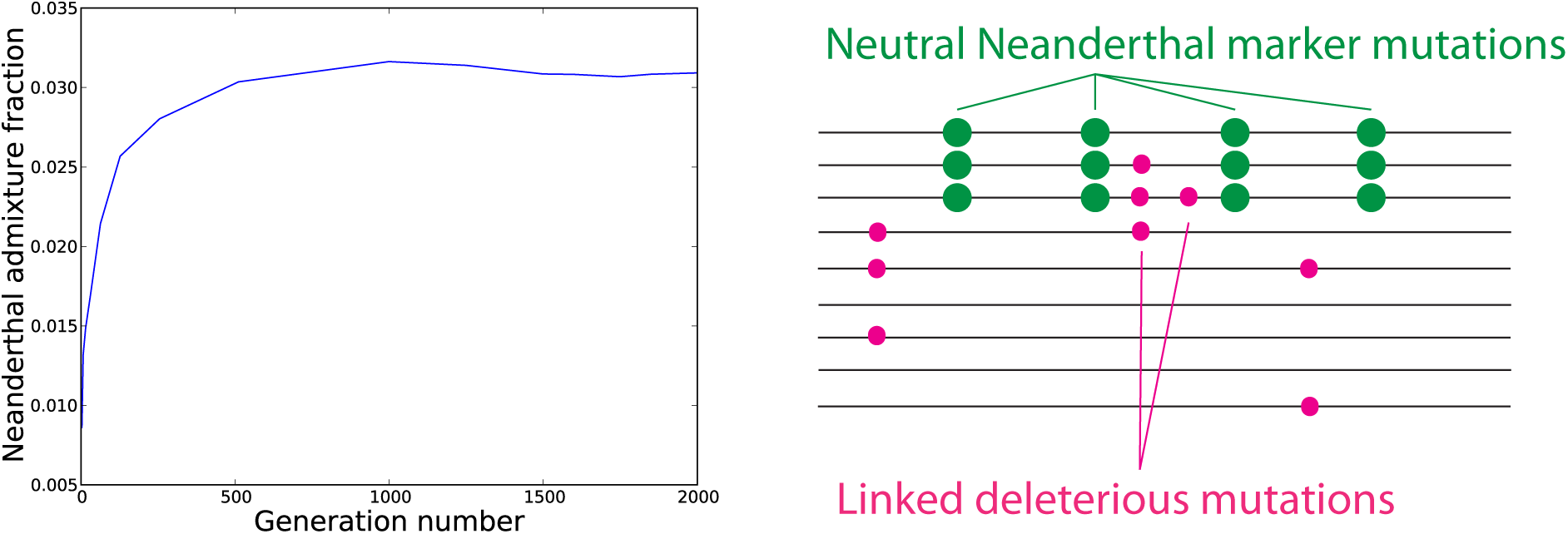
This plot depicts Neanderthal admixture over 2,000 generations in a simulation where all mutations are recessive. The initial admixture fraction is only 1%, but rises to 3% genomewide due to selection for Neanderthal haplotypes that protect against human recessive mutations. The accompanying cartoon illustrates how neutral marker mutations (one every 10^5^ base pairs) are used to measure the Neanderthal admixture fraction as a function of time. The average frequency of these markers, an estimate of the total admixture fraction, is reduced by selection on linked deleterious mutations.

Before admixture, most deleterious alleles are private to either humans or Neanderthals, leading introgressed Neanderthal alleles to be hidden from purifying selection when they are introduced at low frequency. Because Neanderthal haplotypes rarely have deleterious alleles at the same sites that human haplotypes do, they are protective against deleterious human variation, despite the fact that they have a much higher recessive burden than human haplotypes.

It is worth noting that these simulation results assume random mating within Neanderthals and archaic humans; if consanguinity were widespread in either population, this could eliminate much recessive deleterious variation. However, if consanguinity were common in Neanderthals and rare in contemporary humans, a plausible scenario given Neanderthals’ smaller population size, we would still expect to see some positive selection for introgressed Neanderthal DNA. The strength of this selection should depend mostly on standing recessive variation within the human population, which would not be affected by Neanderthal inbreeding.

Several studies have pinpointed archaic genes that appear to be under positive selection in humans [10, 12, 14, 39, 40] because they confer resistance to pathogens or are otherwise strongly favored. Examples of recent adaptive introgression also abound in both animals and plants [41, 42, 43, 44], and our results suggest that heterosis could play a role in facilitating this process. Heterosis should not cause foreign DNA to sweep to fixation, but it might prevent introgressed variants from being lost to genetic drift, thereby increasing the probability of their eventual fixation, particularly if the initial introgression fraction is low.

### 2.3 Additive Fitness Effects Lead to Strong Selection Against Early Hybrids

If most deleterious mutations have additive fitness effects instead of being recessive, different predictions emerge. The reduced fitness of Neanderthals is not hidden, but imposes selection against hybrids in the human population. Such selection against negative deleterious mutations could potentially be offset by positive selection or by associative overdominance due to linked recessive mutations. In the absence of these effects, however, we found that an initial admixture fraction of 10% Neanderthals was necessary to observe a realistic value of 2.5% Neanderthal ancestry after 2,0 generations. Most of the selection against Neanderthal ancestry occurred within the first 20 generations after admixture, at which point the average frequency of the Neanderthal markers had already declined below 3% (Figure 4). During the first 20 generations the variance in admixture fraction between individuals is relatively high, permitting efficient selection against the individuals who have more Neanderthal ancestry than others. However, once all individuals have nearly the same admixture fraction but have retained Neanderthal DNA at different genomic locations, Hill-Robertson interference slows down the purging of foreign deleterious alleles[45, 46, 47]. This suggests that introgression of Neanderthal DNA into humans would have been possible without positive selection, despite the high mutational load, but would require a large initial admixture fraction, perhaps close to 10%.

**Figure 4:**
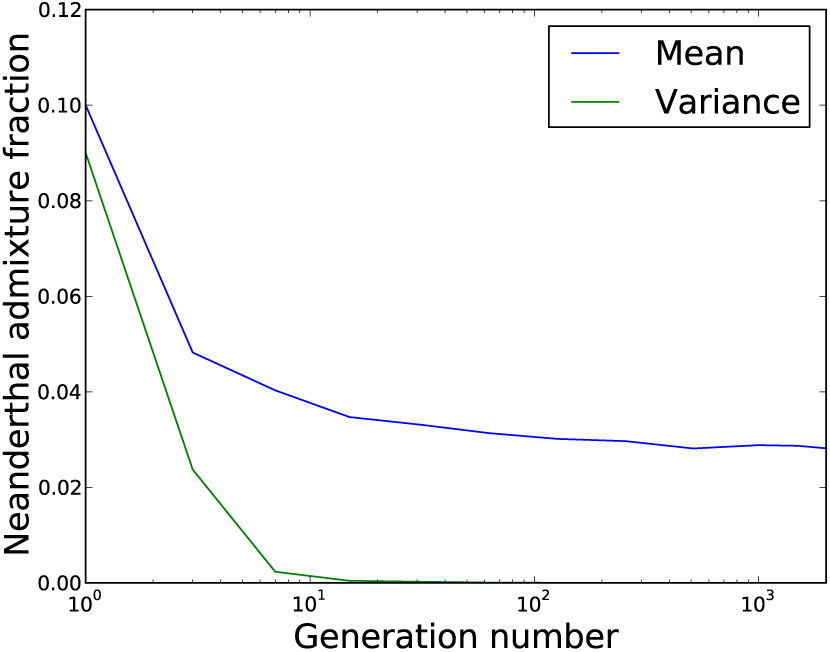
This plot depicts the mean and variance of the Neanderthal ancestry fraction in an admixed population where fitness effects are additive, starting with 10% Neanderthals in generation 1. During the first 10–20 generations after admixture, both the mean and variance in Neanderthal ancestry decrease quickly due to selection against individuals whose Neanderthal ancestry fraction is higher than the population average. After 20 generations, however, all individuals have nearly the same amount of Neanderthal ancestry and selection against its deleterious component becomes less efficient.

Given the qualitative difference between additive and recessive mutation dynamics, we also simulated introgression of partially recessive mutations to see whether they would behave more like additive or fully recessive mutations. In a scenario where all mutations had dominance coefficient *h* = 0.1 and where the initial admixed population contained 10% Neanderthals, we found that partially recessive mutations behaved more like additive mutations than like completely recessive mutations, causing selection against Neanderthal DNA and reduction of the admixture fraction to about 5.5% (Figure 5). This suggests that a significant decline in Neanderthal ancestry with time should be expected under any model where fitness effects are multiplicative across loci and few mutations are completely recessive.

**Figure 5:**
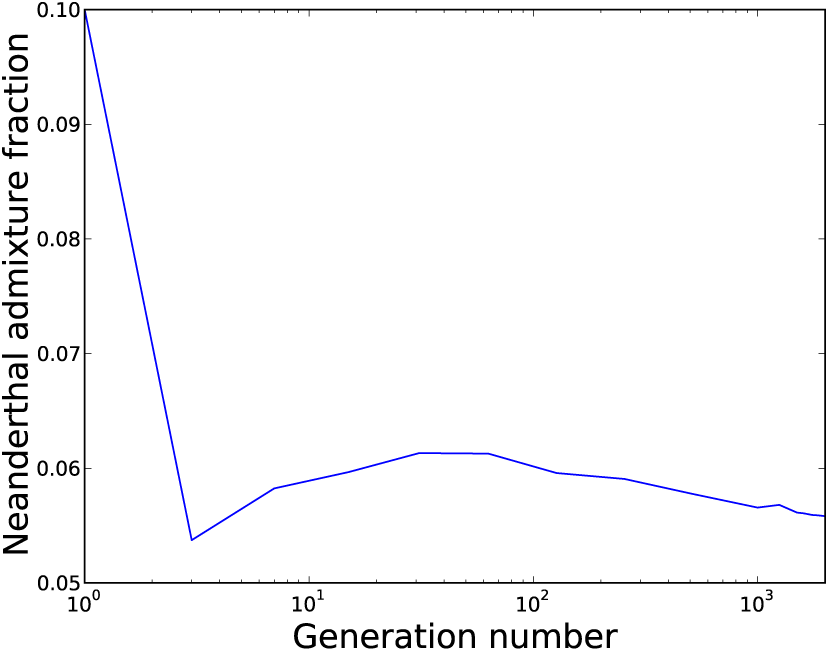
This plot shows how Neanderthal ancestry varies with time in a population where all mutations are partially recessive with dominance coefficient *h* = 0.1. Interestingly, selection against partially recessive foreign alleles was not monotonic; the Neanderthal admixture fraction actually increased due to heterosis for a few tens of generations after undershooting its asymptotic value.

### 2.4 Persistence of Deleterious Neanderthal Alleles in Modern Humans

Figures 1 and 4illustrate two predictions about Neanderthal introgression: first, that it probably introduced many weakly deleterious alleles, and second, that a large fraction of deleterious alleles with additive effects were probably eliminated within a few generations. However, it is not clear from these figures how many deleterious Neanderthal alleles are expected to persist in the present day human gene pool. To address this question, we simulated a control human population experiencing additive mutations that has undergone the out of Africa bottleneck without also experiencing Neanderthal introgression.

At a series of time points between 0 and 2000 generations post-admixture, we recorded each individual’s total load of weakly deleterious mutations (*s* < 0.0005) as well as the total load of strongly deleterious mutations (*s* > 0.0005). The three quartiles of the fitness reduction due to weakly deleterious mutations are plotted in Figure 6A, while the three quartiles of the strongly deleterious fitness reduction are plotted in Figure 6B. Neither the out of Africa bottleneck nor Neanderthal admixture has much effect on the strong load. However, both the bottleneck and admixture exert separate effects on the weak load, each decreasing fitness on the order of 1%.

**Figure 6:**
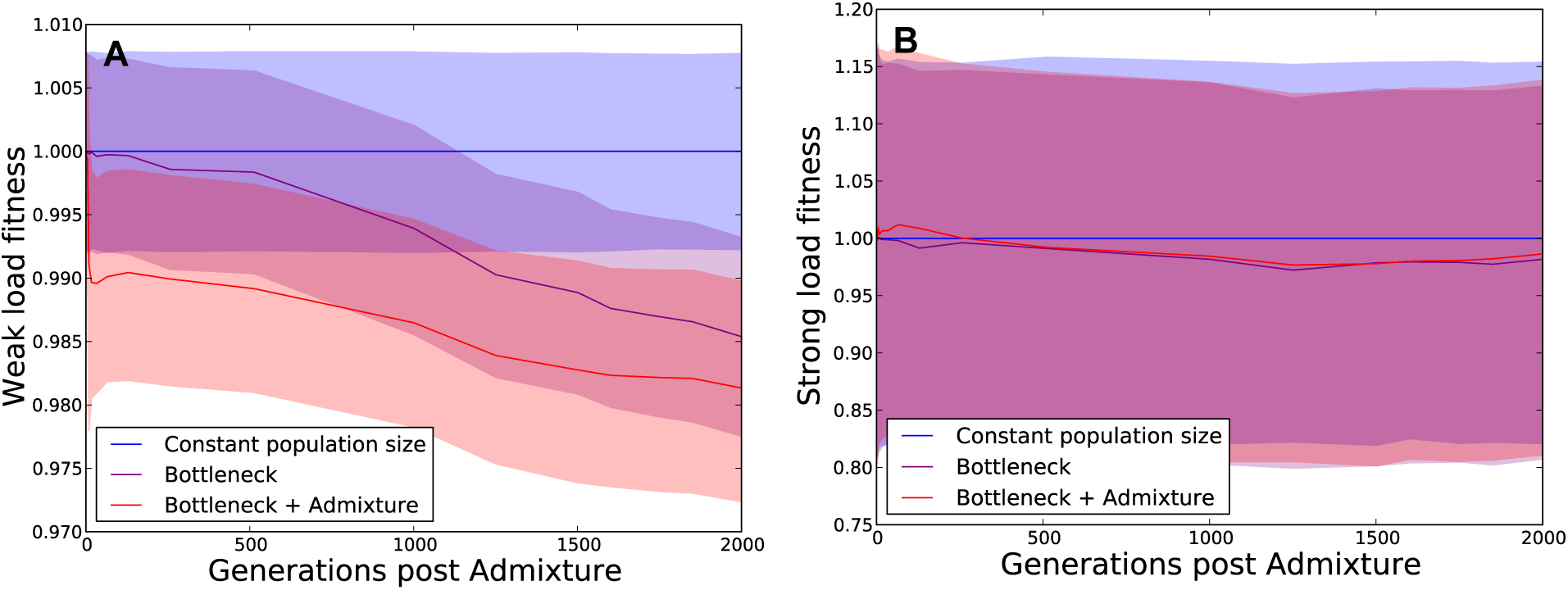
These plots show the mutation load in three simulated human populations, one with constant population size, one that experiences only the out of Africa bottleneck and a third that experiences the bottleneck along with Neanderthal admixture (see Figure 2), assuming additive fitness effects. We partition each individual’s mutation load into two components: the weak load due to mutations with selection coefficient less than 0.0005, and the strong load due to mutations with selection coefficient greater than 0.0005 (note the difference in scale between the two *y* axes). At time *t*, each individual’s weak-load fitness and strong-load fitness are normalized relative to the median individual in the constant-size population. The solid lines show the median in each respective population, while the shaded area encompasses the 25th through 75th percentiles. Panel A shows that the admixed population suffers the greatest fitness reduction due to weak mutations, even 2,000 generations after admixture. Panel B shows that neither the bottleneck nor admixture affects the strong load.

The excess weak load attributable to Neanderthal admixture is much smaller than the variance of strong mutation load that we observe within populations, which is probably why the excess Neanderthal load decreases in magnitude so slowly over time. However, the two load components have very different genetic architectures–the strong load consists of rare variants with large fitness effects, whereas the weak load is enriched for common variants with weak effects. Although surviving Neanderthal alleles are unlikely to affect the risks of Mendelian diseases with severe effects, they may have disproportionately large effects on polygenic traits that influence fitness.

### 2.5 Depletion of Neanderthal Ancestry near Genes can be Explained without Reproductive Incompatibilities

Looking at empirical patterns of human-Neanderthal haplotype sharing, Sankararaman, et al. found that Neanderthal ancestry appears to be depleted from conserved regions of the genome [10]. In particular, they found that the Neanderthal ancestry fraction appears to be negatively correlated with the B statistic, a measure of the strength of background selection as a function of genomic position [48]. In the quintile of the genome that experiences the strongest background selection, they observed a median Neanderthal ancestry fraction around 0.5%, while in the quintile that experiences the weakest background selection, they calculated a median admixture fraction around 2%. This has been interpreted as evidence for epistatic reproductive incompatibilities between humans and Neanderthals [10, 58].

In light of the strong selection against Neanderthal DNA we have predicted on the basis of demography, we posit that reproductive incompatibilities are not required to explain much of the Neanderthal ancestry depletion observed near conserved regions of the genome. Conserved regions are regions where mutations have a high probability of being deleterious and thus being eliminated by natural selection; these are the regions where excess weakly deleterious mutations are most likely to accumulate in Neanderthals. This suggests that selection will act to reduce Neanderthal ancestry in conserved regions even if each allele has the same fitness in both populations.

To model the impact of background selection on patterns of Neanderthal introgression, we explored how the Neanderthal ancestry fraction is expected to decrease over time in the neighborhood of a site where Neanderthals are fixed for a weakly deleterious allele. Using theory and simulations, we show that purifying selection is expected to reduce the frequency of both the deleterious variant and of linked Neanderthal DNA spanning approximately one megabase.

Assuming that Neanderthals are fixed for a deleterious variant that is absent from the human gene pool before introgression, it is straightforward to calculate the expected admixture fraction at a linked neutral locus as a function of time. Given a neutral allele *a* located *L* base pairs away from a deleterious allele of selection coefficient *s*, the frequency *f_a_* of allele *a* is expected to decrease every generation until the deleterious allele recombines onto a neutral genetic background. Letting *r* denote the recombination rate per site per generation and *f_a_*(*T*) denote the frequency of allele *a* at time *T*, then

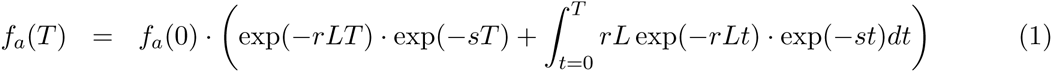

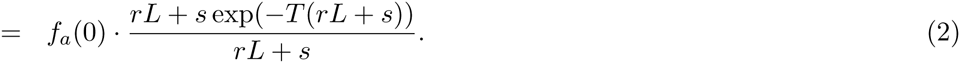

The first term inside the parenthesis of equation (1) is the probability that the two-locus Neanderthal haplotype will remain intact for *T* generations, multiplied by the expected reduction in frequency of the deleterious allele. The integrand of the second term is the probability that this haplotype will instead be broken up by a recombination occurring *t* < *T* generations postadmixture, multiplied by the expected reduction in allele frequency during those *t* generations of linkage. The sum of the constant term and the integral is the expected reduction in frequency of the neutral allele *a*, marginalized over all possible lengths of time that it might remain linked to the deleterious allele.

If the deleterious allele is not fixed in Neanderthals before introgression, but instead has Neanderthal frequency *f_N_* and human frequency *f_H_*, the expected admixture fraction after *T* generations is instead

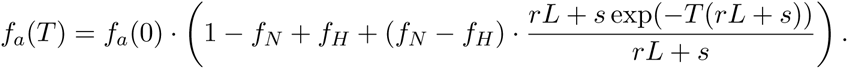

This can be viewed as a case of associative overdominance as described by Ohta, where linked deleterious alleles reduce the expected frequency of a neutral allele down to a threshold frequency that is determined by the recombination distance between the two loci [49].

Figure 7 shows the expected Neanderthal admixture fraction in the neighborhood of a site where Neanderthals are fixed for a deleterious variant of selection coefficient *s* = 5 × 10^−4^. This selection coefficient lies in the middle of the range that is expected to be differentially retained in humans and Neanderthals (see Figure 1). The initial Neanderthal admixture fraction is set at 2%, but after 2,000 generations, the deleterious variant is segregating at only 0.7% frequency on average. Even at a distance of 60 kb from the site under selection, the Neanderthal admixture fraction has declined two-fold from its initial value.

**Figure 7:**
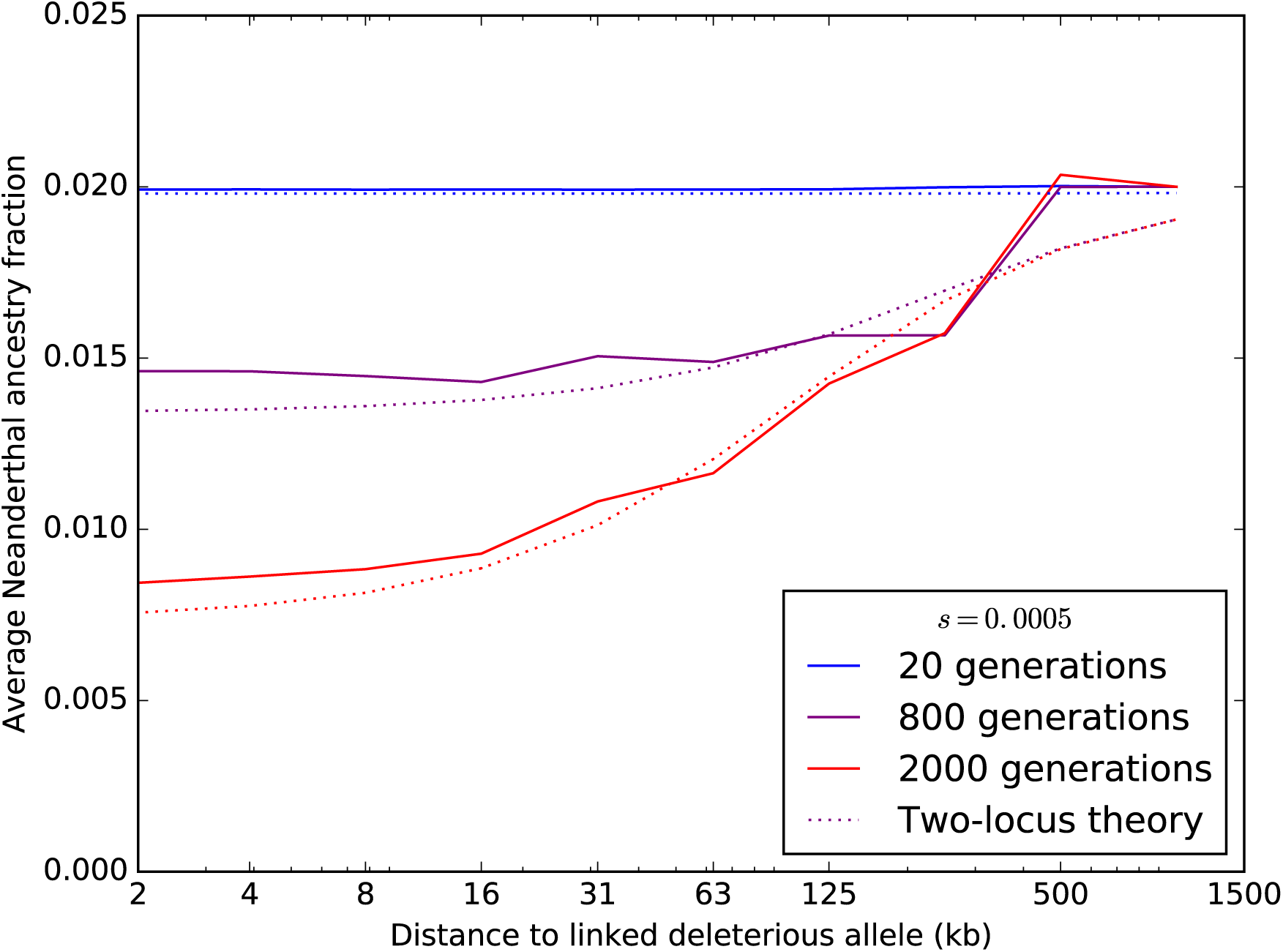
Each colored curve plots the average fraction of Neanderthal ancestry after *T* generations of admixture (*T* = 20, 800, or 2,000) as a function of distance from an introgressed deleterious mutation, assuming an additive model of dominance effects. Solid lines show simulation results averaged over 300 SLiM replicates. For each time point, a dotted line of the same color as the simulation line shows the Neanderthal ancestry fraction profile that is predicted by Equation (2).

We estimated earlier that Neanderthals were approximately 60% as fit as humans, assuming that deleterious mutations have additive fitness effects (see Figure 1A). If this load were composed entirely of variants with selection coefficient *s* = 5 × 10^−4^, this would imply that the typical diploid Neanderthal genome contained – log(0.6)/(5 × 10^−4^) ≈ 1,022 more deleterious variants than the typical diploid human. Distributed across a genome of length 6 × 10^9^ base pairs, this amounts to one deleterious allele every 5.5 × 10^6^ nucleotides. If each deleterious variant causes significant depletion of Neanderthal DNA from a 1-megabase region, this depletion should affect 20% of the genome in a highly noticeable way. Our calculation obviously oversimplifies human genetic architecture, as we do not expect deleterious variants to be evenly spaced or have identical selection coefficients. However, it suggests the archaic mutation load may have been substantial enough to cause background selection against linked neutral DNA across a large proportion of the genome.

## 3 Discussion

Our simulations show that an increased additive mutational load due to low population size is sufficient to explain the paucity of Neanderthal admixture observed around protein coding genes in modern humans. However, our results do not preclude the existence of Dobzhansky-Muller incompatibilities between Neanderthals and humans. Other lines of evidence hinting at such incompatibilities lie beyond the scope of this study. One such line of evidence is the existence of Neanderthal ancestry “deserts” where the admixture fraction appears near zero over stretches of several megabases. Another is the depletion of Neanderthal ancestry near testis-expressed genes [10] and recent chromosomal rearrangements [50]. However, these patterns could be explained by a relatively small number of negative epistatic interactions between human and Neanderthal alleles, as only 10–20 deserts of Neanderthal ancestry have been identified.

Depletion of Neanderthal DNA from the X chromosome has also been cited as evidence for reproductive incompatibilities, perhaps in the form of male sterility [10]. However, we note that the X chromosome may have experienced more selection due to its hemizygous inheritance in males that exposes recessive deleterious mutations [51, 52]. We have shown that selection against the first few generations of hybrids is determined by the load of additive (or hemizygous) mutations, and that the strength of this initial selection determines how much Neanderthal DNA remains long-term. This implies that the admixture fraction on the X chromosome should be lower than on the autosomes if some deleterious mutations are recessive, even in the absence of recessive incompatibility loci that are thought to accumulate on the X according to Haldane’s rule [53, 54, 55]. Frequent selective sweeps also appear to have affected the ampliconic regions of the X chromsome [56], and it is not clear what effect these sweeps may have had on the presence and detection of archaic gene flow.

A model assuming that the general pattern of selection is caused by epistatic effects would involve hundreds or thousands of subtle incompatibilities in order to explain the genome-wide negative correlation of Neanderthal ancestry with background selection. Given the relatively recent divergence between humans and Neanderthals and the abundant evidence for their admixture, it seems unlikely that this divergence could have been given rise to hundreds of incompatible variants distributed throughout the genome. In contrast, our results show that it is highly plausible for the buildup of weakly deleterious alleles to reduce the fitness of hybrid offspring, causing background selection to negatively correlate with admixture fraction.

Neanderthals were not the only inbred archaic population to interbreed with anatomically modern humans. Their sister species, the Denisovans, appears to have contributed DNA to several populations outside Africa, most notably in Oceania and Asia [57, 58, 59]. Since genetic diversity appears to have been comparably low in Denisovans and Neanderthals, owing to a bottleneck of similar duration and intensity [57, 2], our inferences about the action of selection on Neanderthal DNA should apply to Denisovan DNA with equal validity. Like Neanderthal introgression, Denisovan DNA appears to be depleted near genes and also depleted from the X chromosome relative to the autosomes [58].

The distribution of dominance effects in humans is not well characterized. But it is likely that introgressed Neanderthal DNA has been subject to a selective tug-of-war, with selection favoring Neanderthal DNA in regions where humans carry recessive deleterious mutations and selection disfavoring Neanderthal alleles that have additive or dominant effects. In a sense, this is the opposite of the tug-of-war that may occur when a beneficial allele is linked to recessive deleterious alleles that impede the haplotype from sweeping to high frequencies [60, 61].

If most mutations have fitness effects that are additive and multiplicative across loci, initial admixture would have to have been as high as 10% to explain the amount of admixture observed today. In contrast, if most mutations are recessive, an initial admixture fraction closer to 1% appears most plausible. Large changes in admixture fraction are predicted as a consequence of strong deleterious effects that result from long-range linkage among hundreds of weakly deleterious alleles. Kim and Lohmueller previously simulated introgression scenarios that included a wider range of selection and dominance coefficients, but as their simulations did not include linkage, they found that more strongly deleterious alleles were required to change admixture proportions over time [62]. Likewise, Juric, et al. modeled linkage among Neanderthal alleles at a megabase scale but not a genome-wide scale and observed not much change in admixture fraction over time [63]. They did, however, infer depletion of Neanderthal DNA near genes, concluding independently of our work that this observation could be a consequence of the Neanderthal population bottleneck.

We did not model selection for new beneficial mutations here, and it is possible that such selection might also have helped facilitate introgression, particularly in the first generations where selection against hybrids would otherwise have been strong. As more paleolithic human DNA is sequenced, it may become possible to measure how admixture has changed over time and extract information from this time series about the distribution of dominance coefficients. This information could also help resolve confusion about the fitness effects of the out of Africa bottleneck, which is predicted to have differently affected the burdens of additive versus recessive variants [17, 33].

We do not claim to have precisely estimated the deleterious Neanderthal load that remains in non-Africans today, as this would require better estimates of the DFE across different genes and more exploration of the effects of assumptions regarding recent demographic history. However, our results suggest that Neanderthal admixture should be incorporated into models exploring mutational load in humans to more accurately predict the mutation load difference between Africans and non-Africans. Association methods have already revealed correlations between Neanderthal alleles and several human diseases [10]. Our results on mutations with additive dominance effects suggest that introgression reduced non-African fitness about as much as the out-of-Africa bottleneck did.

Introgression of recessive mutations is predicted to affect fitness in a more complex way. Some adaptive benefits will result from Neanderthal and human haplotypes masking one another’s deleterious alleles, but Hill-Robertson interference may also hurt fitness as overdominant selection at recessive sites drags linked dominant Neanderthal alleles to higher frequency. In addition, Neanderthal haplotypes are predicted to have worse recessive burdens than human ones if they become homozygous due to selection or inbreeding.

Our results have implications for conservation biology as well as for human evolution, as they apply to any case of secondary contact between species with different effective population sizes.

When an outbred population experiences gene flow from a more inbred population, we predict an increase in genetic entropy where deleterious alleles spill rapidly into the outbred population and then take a long time to be purged away by selection. This process could magnify the effects of outbreeding depression caused by genetic incompatibilities [64, 65, 66] and acts inversely to the genetic rescue process, in which individuals from an outbred population are artificially transplanted into a threatened population that has been suffering from inbreeding depression [67, 68, 69]. These results suggest that care should be taken to prevent two-way gene flow when genetic rescue is being attempted to prevent lasting damage to the fitness of the outbred population.

## 4 Acknowledgements

We thank Joshua Schraiber and Benjamin Vernot for manuscript comments and members of the Nielsen, Slatkin, and Pritchard labs for helpful discussions. K.H. received support from a Ruth L. Kirschstein National Research Service Award from the National Institutes of Health (award number F32GM116381). K.H. and R.N. also received support from NIH Grant IR01GM109454-01 to R.N., Yun S. Song, and Steven N. Evans. The content of this publication is solely the responsibility of the authors and does not necessarily represent the official views of the National Institutes of Health.

